# A revisit of RSEM generative model and its EM algorithm for quantifying transcript abundances

**DOI:** 10.1101/503672

**Authors:** Hy Vuong, Thao Truong, Thang Tran, Son Pham

## Abstract

RSEM has been mainly known for its accuracy in transcript abundance quantification. However, its quantification time is extremely high compared to that of recent quantification tools. In this paper, we revised the RSEM’s EM algorithm. In particular, we derived accurate M-step updates to eliminate incorrect heuristic updates in RSEM. We also implement some optimizations that reduce the quantification time about a hundred times while still have better accuracy compared to RSEM. In particular, we noticed that different parameters have different convergence rates, therefore we identified and removed early converged parameters to significantly reduce the model complexity in further iterations, and we also use SQUAREM method to further speed up the convergence rate. We implemented these revisions in a packaged named Hera-EM, with source code available at: https://github.com/bioturing/hera/tree/master/hera-EM

## 1 Introduction

RNA-Seq has become a standard approach for quantifying transcript abundances. Reads with multiple mapping locations (multi-mapping reads) are the main obstacle in quantifying. Many tools have been developed to resolve this problem. The simplest solution is ignoring multi-mapping reads, and just counting the number of unique reads for each transcript. This approach has limitations [6]. To take into account information from multi-mapping reads, EM solutions were developed [2, 6,10, 11]. Using a comprehensive generative model, RSEM [6] has been known for its accuracy in transcript abundance quantification [2, 9]. However, given the large amount of RNA-Seq data generated, the running time of RSEM is a major disadvantage. This gives rise to a new class of more efficient tools e.g. Sailfish, Salmon, Kallisto [2, 10, 11]. Besides using pseudo-alignment instead of alignment. Kallisto, and Sailfish implement simpler EM solutions that group reads mapping to the same set of transcript into equivalent classes^1^ which may lead to loss of mapping positions, fragment lengths, alignment score information.^2^. In the case of multi-mapping reads, these variables are important to find the best assignments of reads to transcripts. As a result, Kallisto and Sailfish, while being much faster, cannot reach the accuracy level of RSEM [2,9].

In the work of Li et. al., 2011 [6], while the generative RSEM model was constructed comprehensively, the authors failed to derive accurate update formulae for the models’ parameters in their EM algorithm as the difficulty of the analytic work. In this manuscript, we keep the essence of RSEM’s model, and provide accurate formulae for parameter updates in the EM algorithm by finding the solutions analytically. With the accurate parameter update formulae, we achieve better results than RSEM in the SMC DREAM challenge data sets. We note that the data sets from the SMC DREAM challenge were generated using the RSEM generative model.

Besides, we observed that different transcripts have different convergence rates for their abundance estimations. Therefore, by avoiding the recalculation of early converged estimations, we can save a huge amount of computation. Specifically, if we detect a parameter unchanged for a few EM rounds, we will keep the parameter fixed in further iterations. Hence, we can skip the recalculation of the joint probability on those transcripts. This is our main contribution in speeding up the EM process. In addition, we also implement the SQUAREM method [16] to improve the convergence rate.

Hera-EM is about a hundred times faster, at the same time, gains equal or even better accuracy compared to the EM procedure in RSEM. On a data set with 60 million of reads, RSEM takes about an hour (3432 seconds) for EM step only, while Hera-EM needs half a minute (24 seconds); for an other data set with 75 million of reads, RSEM takes about 1.5 hours (5044 seconds), while Hera-EM takes 39 seconds. Hera-EM consumes about 7 GB of memory in all of these benchmarks.

## 2 Method

### Notations

Let 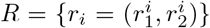 be a set of *N* read pairs; *T* = {*t_i_*} be a set of *M* transcripts. Transcript *t_i_* has length *l_i_*. For each read *r* = (*r*_1_,*r*_2_), we denote *A_r_* = {*a_i_*} as the set of alignments, where each alignment *a_i_* is a tuple (*t, f, s, o*) of transcript *t*, fragment length *f*, starting position *s*, and orientation *o*. We also denote *t_a_*, *f_a_*, *s_a_*, *o_a_* as the original transcript, fragment length, starting position and orientation of an alignment *a*, respectively.

### Generative Model

The process of generating a read is described in the following generative process.

- Draw transcript *t_i_* from the set of all transcript *T* with probability *θ_i_*.
- Choose a fragment length *f_i_* of the read with probability depends on the the universal fragment length parameter *λ* adjusted by the length of transcript *t_i_*.
- Choose a starting position *s_i_* on transcript *t_i_* of the fragment with a universal probability *π* adjusted based on the length of transcript *t_i_*, and the fragment length *f_i_*.
- Choose an orientation *o_i_* of the read pair with probability *ω* of being forward-stranded.
- Adding sequencing errors, with each position has the probability *∊* of being erroneously sequenced.

The details of the above parameters *θ*, *λ*, *π*, *ω*, *∊* are explained below.

### Definition of the distributions and parameters

#### *θ*: Transcript distribution

For transcript distribution, we denote *θ_t_* as the probability that a transcript *t* is chosen from all transcripts to generate a fragment.

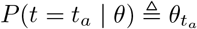

#### *λ*: Fragment length distribution

Given parameter *λ* as the distribution of the fragment length generated from a hypothetical infinitely long transcript. In the generative model of fragments on a particular transcript, this parameter needs to be adjusted according to the length of the transcript, as fragments cannot be longer the transcript itself. We set the probability of the all fragments that are longer than the transcript to zero and normalize the total probability to 1.

Denote *l* = |*t*| as the length of the chosen transcript *t.*

The probability of choosing a fragment length *f_a_*:

If 0 ≤ *f_a_* ≤ *l*

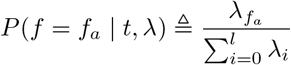

Otherwise

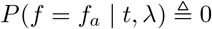

#### *π*: Read start position distribution

As many studies have observed that reads are not generated uniformly along the transcripts [4,6,12], Li and Dewey [6] modeled the read start distribution by dividing each transcript into 20 bins and count the frequency of reads starting in each bin. We also model this distribution by dividing each transcript into a fixed number of segments (64 by default) and assign a common probability *π_i_* to the *i*^th^ segment of every transcript. Given a hypothetical fragment of length 1, *π_i_* represents the probability of this fragment originated from the segment *i.* Within each segment, the starting positions of the fragments are uniformly distributed. With fragment lengths larger than 1, not every starting position on the transcript is possible because fragments can not end outside the transcript, so we need to adjust the distribution.

Let *π_i_* be the probability that a hypothetical fragment of length 1 is originated from the segment *i.* Denote *b* as the number of segments of each transcript, *l* as the length of the chosen transcript.

We define *g*(*k*) as the cumulative distribution function for the hypothetical fragment of length 1.

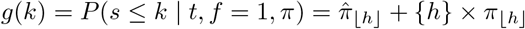

Where 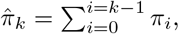 and 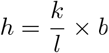.

With a fragment of length *f* originated from transcript *t* of length *l*, the fragment can only start from 0 ≤ *s_a_* ≤ *l* – *f*, so we have the following definition for 0 ≤ *s_a_* < *l* – *f*,

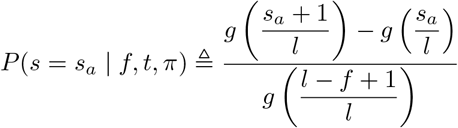

Otherwise,

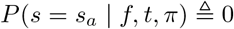

#### *ω*: Orientation distribution

We assume the distribution of the read orientation *o* is a Bernoulli distribution with parameter *ω*. *o* = 1 means the read is forward-stranded, and *o* = 0 means the read is reverse-stranded. We can set 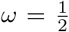 in unstranded protocol, and *ω* = 1 in stranded protocol, or by default it is estimated from the data.

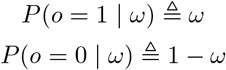

#### *∊*: Error rate

Sequencing error also needs to be accounted for. We assume a simple distribution, every base has the same probability for being erroneously sequenced^3^, denoted as *∊*. Therefore a read *r* with edit distance *e* to the reference has the probability of *∊^e^* × (1 – *∊*)^|*r*|–*e*^.

Given an alignment *a*(*t*, *f*, *s*, *o*) of the read *r*(*r*_1_, *r*_2_), let *d* be the edit distance between the read and the reference of the aligned position.

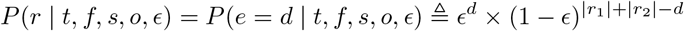

### Likelihood function

Denote Ω as the set of all parameters *θ*, *λ*, *π*, *ω*, and *∊*. From Fig. 1, we derive a formula for joint probability of read *r* and alignment *a*(*t, f, s, o*) given parameter Ω.

**Figure 1:**
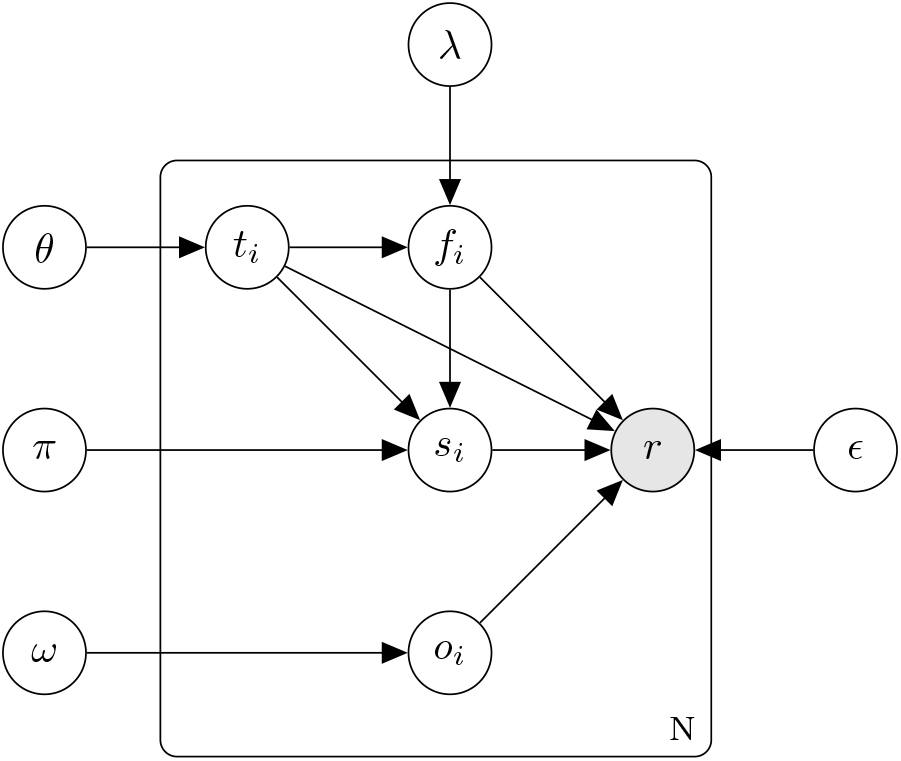
Bayesian graph for read generating process. *r* = (*r*_1_,*r*_2_) is the only observed variable; *t_i_, f_i_, s_i_, o_i_* are the latent variables; *θ*, *λ*, *π*, *ω*, and *∊* are the parameters

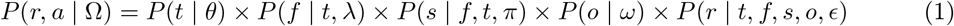

We also have the conditional probability,

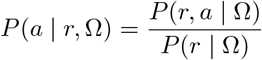

With *P*(*r* | Ω) = Σ_*a∊A_r_*_ *P*(*r, a* | Ω)

We define the likelihood function.

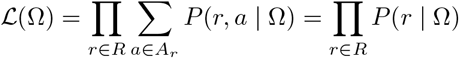

Where *A_r_* is the set of all alignments of a read pair *r*. *R* is the set of all read pairs.

Our goal is to find the parameter Ω that maximizes the likelihood function *𝓛*. In this work, we use an optimized EM algorithm to obtain the desired parameters. First we initialize our distributions as uniform distributions, and then applying the following EM update iteratively until the relative change of all elements in the parameter *θ* is below 10^-3^

### EM update

For each iteration of the EM algorithm, from the old parameter Ω^(*i*)^, we need to find a new parameter Ω^(*i*+1)^ that maximizes this function.

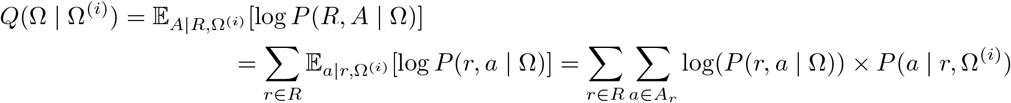

Where *A* is the set of alignments of R.

Taking partial derivatives of *Q*(Ω | Ω^(*i*)^) by the following variables *θ*, *λ*, *π*, *ω*, *∊* and equating them to zero, we can solve for the roots of those equations with the constraints that the sum of each variable equals to 1 (See supplementary for detailed calculations). This gives us the following update formulae.

#### Transcript distribution

For each transcript *t*, we have the following update,

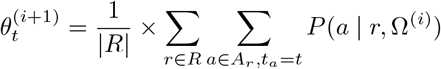

#### Fragment length distribution

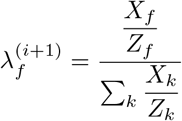

Where *X_f_* = Σ_*r*∊*R*_Σ_*a*∊*A*_*r*_,*f_a_*=*f*_ *P*(*a* | *r*,Ω^(*i*)^), *Y_f_* = Σ_*r*∊*R*_ Σ_*a*∊*A*_*r*_,*f*=|*t_a_*|_ *P*(*a* | *r*,Ω^(*i*)^), *Z*_0_ = 1, and 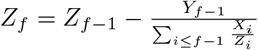

#### Read start position distribution

Let 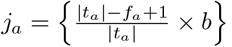, and 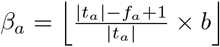

Let *X_k_* = Σ_*r*∊*R*_ Σ_*a*∊*A_r_*,*q_a_*=k_ *P*(*a* | *r*, Ω^(*i*)^)

Let 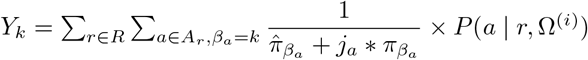

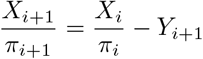

Let 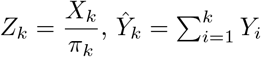

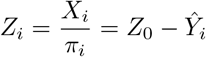

We cannot eliminate the variable *π_i_* in 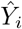 so we use an approximation from the previous value of *π_i_*. We can easily compute the estimation of *Y_i_* alongside with the EM algorithm. However there is still an unsolved variable *Z*_0_, we use the Newton method [1] to solve the equation 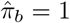 to get the value of *Z*_0_

After finding *Z*_0_, we have

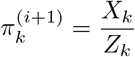

#### Orientation distribution

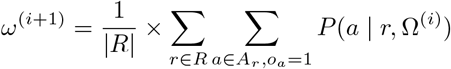

#### Error rate

We have the following update formula,

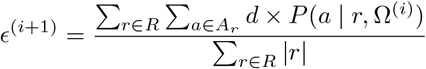

#### Inaccuracy in RSEM parameter updates

While our generative model is the same as RSEM, the update formulae for the parameters *π* and *λ* are different. RSEM does not solve the partial derivative equations to derive the formulae for these two parameters, but rather uses intuition to come up with incorrect update formulae.

In particular, in the RSEM published source code, the parameter *λ*^(*i*+1)^ is updated as follow, 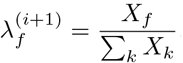, which is not the optimal result for this optimization. Similar to fragment length distribution, RSEM uses an intuitive update formula 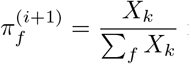 for the parameter *π* of read start position distribution. This leads to underestimating read coverage in the 3’ tails. The error in the update of *read start position* distribution affects the results significantly. In the Results section, we show that RSEM incorrectly estimates the read start distribution, and leads to lower correlations with the simulated truth.

### Optimizations

As the number of reads is extremely high in each run, it yields a heavy computational cost in each naive EM iteration. Many recent methods try to reduce the complexity by grouping similar reads^4^ into equivalent classes [2,10,11]. This approach significantly speeds up the computation, however, with a notably loss in accuracy. In Hera-EM, our goal is to preserve the accuracy, while reducing the quantification time.

Similar to RSEM, we only run full EM iterations for a few first iterations (default of 30 in our implementation), after that we run simplified EM iterations, which only *θ* is being updated, other parameters are kept fixed. This saves a lot of computation as other parameters tend to converge after only a few rounds of EM. Besides, we only need to keep the edit distance for each alignment instead of the whole read sequence to calculate the joint probability. Therefore, we can store the whole alignment data on RAM with even less memory than RSEM. Hence, we don’t have to reread the alignment file in every EM round. This also significantly speeds up each EM iteration.

An interesting observation is that after just a few EM rounds, only a small number of elements in parameter *θ* are still being changed as most of them have already been converged early. This leads us to a more sophisticated optimization. If we detect a parameter *θ_i_* of a transcript have a relative change smaller than a threshold of 10^-5^ in a few EM rounds, the parameter will be kept fixed in further iterations. This means the joint probability of the alignments on these transcripts will not change in further iterations and we don’t need to update the parameters *θ_i_* of those transcript, therefore they can be safely removed from the future iterations. We only need to keep the total joint probability of the removed alignments for each read to calculate the conditional probability of other alignments. Applying this optimization, we see a dramatic reduction of data complexity. This is our main optimization of the EM algorithm.

We noticed that TIGAR2 [7] utilizes a similar optimization, which stops the re-evaluation of reads that mapped onto the transcripts with converged parameters. However, there is a notably difference between TIGAR2 and our method. In our method, we don’t wait until all alignments of a read to be converged to stop re-evaluating that read. We stop the re-evaluating of an alignment immediately after that alignment is converged, independent from other alignments of the same read.

We also ignore the improbable alignments with conditional probability *P*(*a* | *r*, Ω) less than a defined threshold. Through experimenting, we notice that while a smaller threshold of 10^-6^ give a nearly identical result compared to no-filtered version, higher threshold of 10^-3^ is notably faster but can cost accuracy. Hence we choose a default of 10^-6^. In our simulated data sets, about half of the alignments are redundant due to very low conditional probability, therefore, this optimization reduces about half of the quantification time.

Finally, we also implement SQUAREM [16] to speed up the convergence rate.

### TPM value

Similar to RSEM [6], we defined the effective length of the transcript 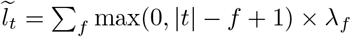.

The TPM value of a transcript *t* is 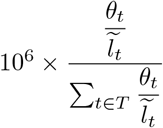

## 3 Results

We benchmarked Hera-EM algorithm against RSEM using two simulated data sets generated by SMC DREAM challenge [14]. The first data set denoted as *sim*4*x* contains 5 samples: 41, 42, 43, 44, 45; each sample has 60 million reads. The second data set, denoted as *sim*5*x*, is an improved version of *sim*4*x*, and contains 6 samples: 51, 52, 53, 54, 55, 56; each sample has 75 million reads. Both data sets are simulated by rsem-simulate-reads. The main difference between *sim*5*x* and *sim*4*x* is the read start position distribution. In *sim*4*x*, reads are uniformly generated along the transcripts, however in *sim*5*x* many biases are introduced to better reflect the situations in real RNA-sequencing data [14,15].

The benchmark was run on a computer with 40-cores Intel(R) Xeon(R) E5-2680 v2 @ 2.80GHz CPU, and 96 GiB DDR3 1866 MHz RAM. Each tool was run with its default parameters in 32 threads. Besides, we also ran RSEM with --estimate-rspd option to test the *read start position* update of RSEM. Hera-EM and RSEM used the same alignment files performed by bowtie2 [5] with the following parameter which is from RSEM.

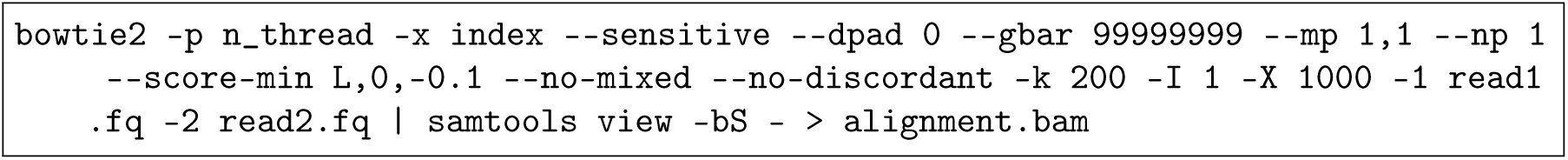

Table 1 describes the running time and peak memory for EM procedures of Hera-EM and RSEM. On average, Hera-EM is about a hundred times faster than RSEM’s EM. Hera-EM consumes about 6 to 8 GBs memory while RSEM EM procedure consumes about 6 to 9 GBs. This table also presents the correlation between both tools’ TPM estimation and the simulated truth TPM. The estimation and the simulated truth TPM values are both rounded to 2 decimal digits before computing the correlation. Spearman’s rank correlation and pearson correlation are computed using their definitions [3]. To compute the log-pearson score, each TPM value is first added an offset of 0.01 then we compute the pearson correlation on the log-transformed values of the modified TPM values (These benchmark metrics are based on the SMC DREAM challenge [13]).

In *sim*4*x* data sets, the results from Hera-EM and RSEM are extremely similar. However, in *sim*5*x* data sets, Hera-EM is more accurate than RSEM in all metrics (Spearman, Pearson, Log-pearson correlations with the ground simulated truth). A scatter plot between the estimation of both tools and simulated truth of the sim 51 is shown in Fig. 2. This difference is because the read start distribution in sim5x is not uniformed, while RSEM assumes it to be uniformed by default.

**Table 1:** Running time, peak memory and correlations with simulated truth TPM of Hera-EM and RSEM

| Dataset | Time (second) |  | Memory (MB) |  | Spearman |  | Pearson |  | Log-Pearson |  |
| --- | --- | --- | --- | --- | --- | --- | --- | --- | --- | --- |
|  | Hera-EM | RSEM | Hera-EM | RSEM | Hera-EM | RSEM | Hera-EM | RSEM | Hera-EM | RSEM |
| sim41 | **32** | 3311 | **6404** | 6804 | 0.9408 | **0.9417** | **0.9988** | 0.9986 | 0.9763 | **0.9766** |
| sim42 | **39** | 2971 | **6388** | 6752 | 0.9375 | **0.9377** | **0.9952** | 0.9941 | 0.9766 | **0.9767** |
| sim43 | **47** | 4035 | **6454** | 6811 | 0.9416 | **0.9425** | 0.9959 | 0.9959 | 0.9773 | **0.9776** |
| sim44 | **42** | 3567 | **6565** | 7080 | 0.9378 | **0.9379** | 0.9995 | 0.9995 | 0.9758 | 0.9758 |
| sim45 | **24** | 3432 | **6384** | 6469 | 0.9379 | **0.9385** | 0.9989 | 0.9989 | 0.9751 | **0.9755** |
| sim51 | **58** | 4290 | **7675** | 8495 | **0.9421** | 0.9291 | **0.9984** | 0.9970 | **0.9761** | 0.9593 |
| sim52 | **74** | 4637 | **7747** | 8614 | **0.9448** | 0.9318 | **0.9994** | 0.9974 | **0.9774** | 0.9610 |
| sim53 | **39** | 5044 | **7778** | 8970 | **0.9433** | 0.9322 | **0.9957** | 0.9956 | **0.9769** | 0.9611 |
| sim54 | **51** | 4514 | **8206** | 9849 | **0.9426** | 0.9293 | **0.9988** | 0.9979 | **0.9755** | 0.9590 |
| sim55 | **48** | 4157 | **7632** | 8496 | **0.9389** | 0.9266 | **0.9998** | 0.9986 | **0.9756** | 0.9599 |
| sim56 | **45** | 4074 | **8083** | 9245 | **0.9373** | 0.9254 | **0.9997** | 0.9890 | **0.9740** | 0.9581 |

**Table 2:** Correlations of Kallisto, Hera-EM, and RSEM with rspd estimation option turned on with simulated truth TPM. Kallisto and Hera-EM were run with default parameters.

| Dataset | Kallisto |  |  | Hera-EM |  |  | RSEM with rspd estimation |  |  |
| --- | --- | --- | --- | --- | --- | --- | --- | --- | --- |
|  | Spearman | Pearson | Log-Pearson | Spearman | Pearson | Log-Pearson | Spearman | Pearson | Log-Pearson |
| sim41 | 0.9057 | 0.9769 | 0.9624 | **0.9408** | **0.9988** | **0.9763** | 0.9332 | 0.9977 | 0.9679 |
| sim42 | 0.9073 | 0.9887 | 0.9647 | **0.9375** | **0.9952** | **0.9766** | 0.9295 | 0.9939 | 0.9679 |
| sim43 | 0.9072 | 0.9928 | 0.9647 | **0.9416** | **0.9959** | **0.9773** | 0.9316 | 0.9958 | 0.9665 |
| sim44 | 0.9050 | 0.9805 | 0.9631 | **0.9378** | **0.9995** | **0.9758** | 0.9309 | 0.9993 | 0.9677 |
| sim45 | 0.9043 | 0.9916 | 0.9621 | **0.9379** | **0.9989** | **0.9751** | 0.9299 | 0.9971 | 0.9670 |
| sim51 | 0.8901 | 0.9881 | 0.9454 | **0.9421** | **0.9984** | **0.9761** | 0.9304 | 0.9973 | 0.9617 |
| sim52 | 0.8922 | 0.9878 | 0.9477 | **0.9448** | **0.9994** | **0.9774** | 0.9310 | 0.9984 | 0.9620 |
| sim53 | 0.8939 | 0.9762 | 0.9479 | **0.9433** | 0.9957 | **0.9769** | 0.9305 | **0.9958** | 0.9609 |
| sim54 | 0.8908 | 0.9908 | 0.9460 | **0.9426** | **0.9988** | **0.9755** | 0.9277 | 0.9979 | 0.9586 |
| sim55 | 0.8938 | 0.9948 | 0.9480 | **0.9389** | **0.9998** | **0.9756** | 0.9243 | 0.9991 | 0.9588 |
| sim56 | 0.8938 | 0.9845 | 0.9458 | **0.9373** | **0.9997** | **0.9740** | 0.9248 | 0.9972 | 0.9567 |

We further run RSEM with --estimate-rspd option to enable the estimation of read start position distribution. As the update formula for this parameter in RSEM is not correct, RSEM’s results deteriorate (Table 2 and Fig.2).

We also benchmarked the accuracy of Kallisto (Table 2) on these data sets. Kallisto has the lowest scores in all metrics. Note that as Kallisto doesn’t provide user the ability to input alignment files, the low accuracy can also be the result of spurious pseudo-alignment process.

We also noticed that given the same bowtie2 alignment, Salmon’s quantification results exceed both RSEM and Hera-EM in the *sim*4*x*, and stay between Hera-EM (better) and RSEM in *sim*5*x*. We believe this may stem from the differences between variational bayesian and EM algorithms.

**Fig. 2:**
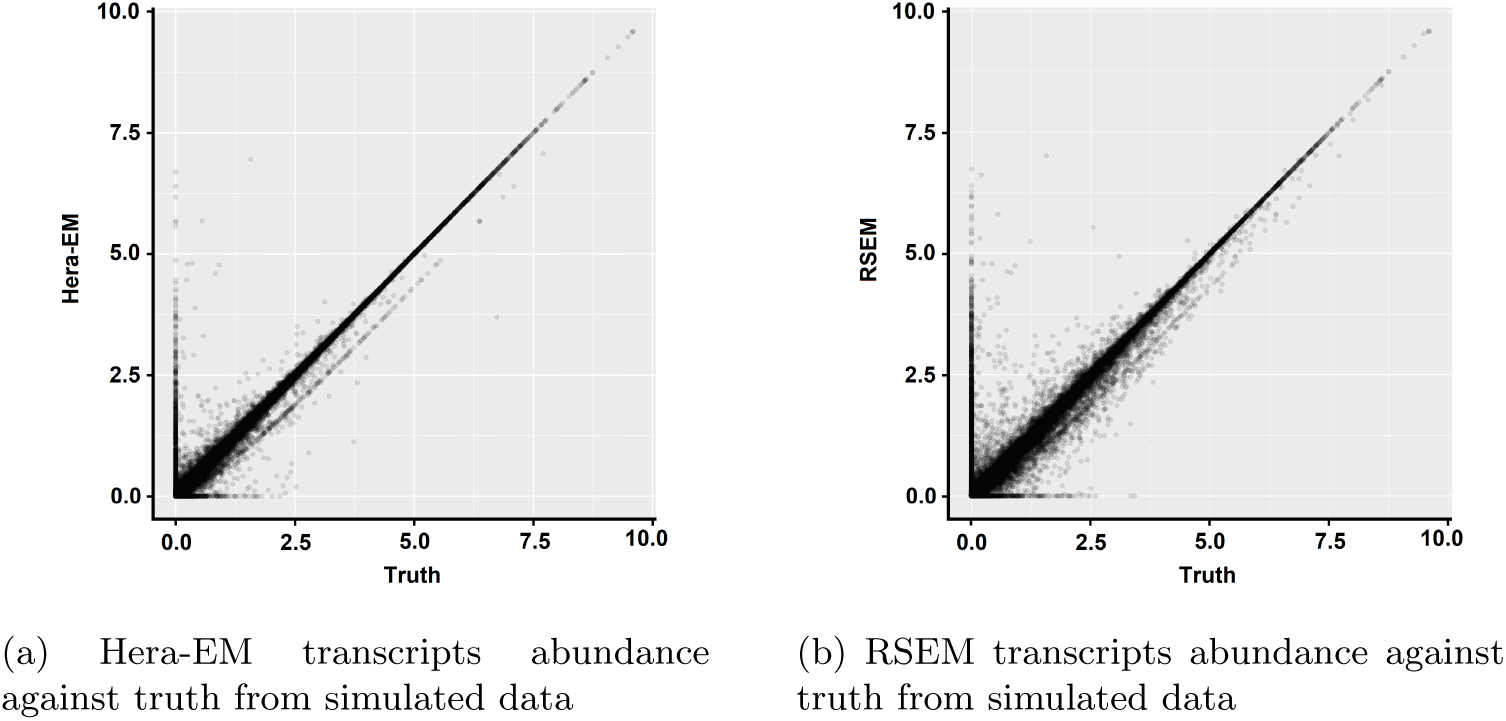
Scatter plots of estimated abundances and truth TPM in sim51 of both tools. Hera-EM shows a better correlation with the simulated truth TPM. All tools were run with default parameters.

## 4 Supplementary

As stated, our goal is to find the parameter Ω to maximize the likelihood *𝓛* using EM algorithm

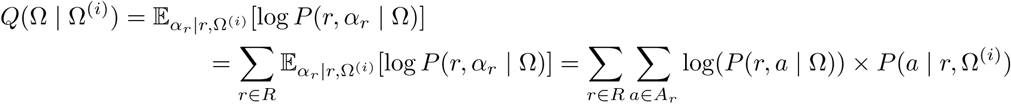

We have to maximize this function with respect to Ω

### 4.1 Transcript distribution

Our goal is to find the maximum value of *Q*(Ω | Ω^(*i*)^) respect to *θ* with the condition ∑_*i*_ *θ_i_* = 1 From the equation 1, we have,

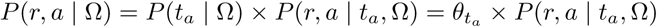

Whereas the probability *P*(*r,a* | *t_a_*,Ω) is independent from *θ*. Hence we have,

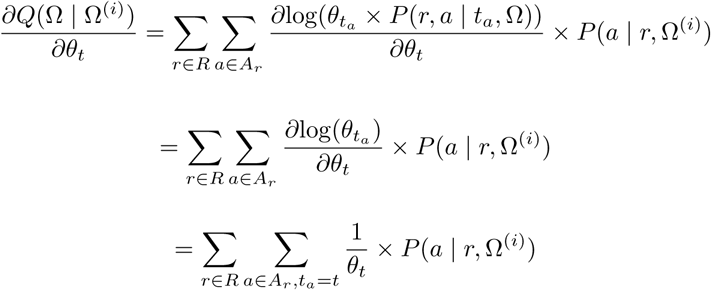

Using Lagrange multiplier, we have the following system of equations,

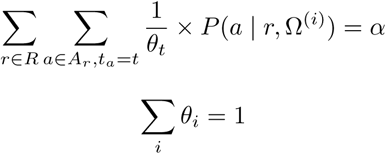

With *α* is the Lagrange multiplier, hence,

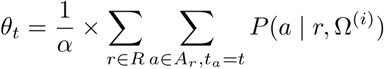

Solve for *α*

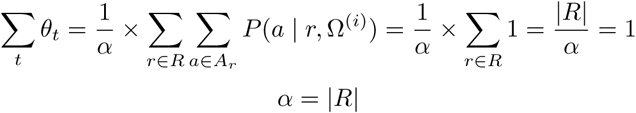

Therefore,

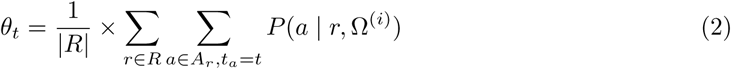

### 4.2 Fragment length distribution

Similarly to transcript distribution, we have,

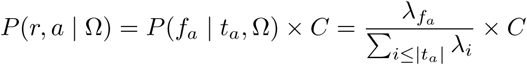

With *C* is some function that is independent from λ Hence,

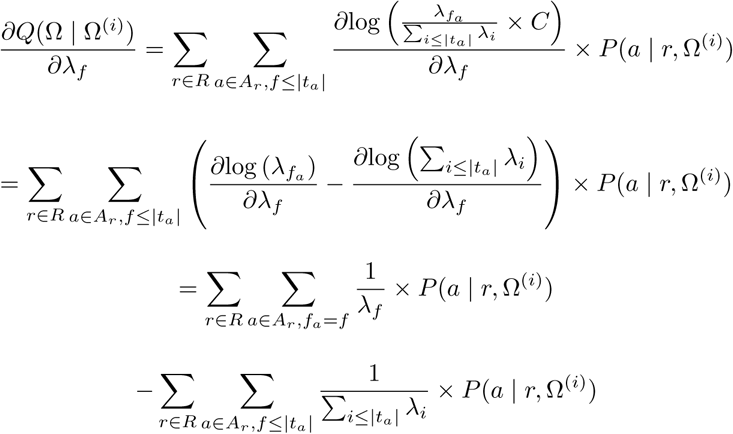

Using Lagrange multiplier, we have the following system of equations,

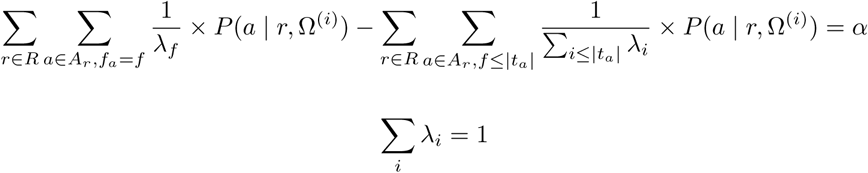

We calculate the difference between the equation for λ_*f*_ and λ_*f*−1_

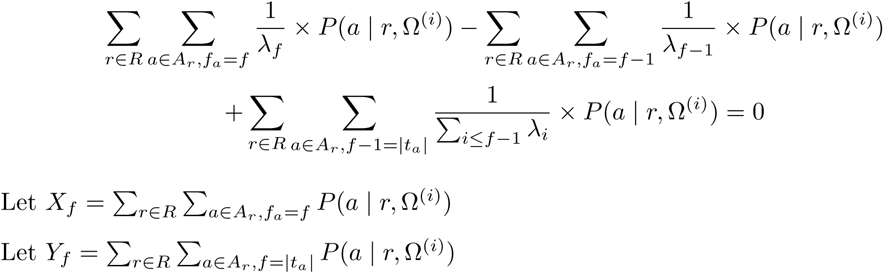

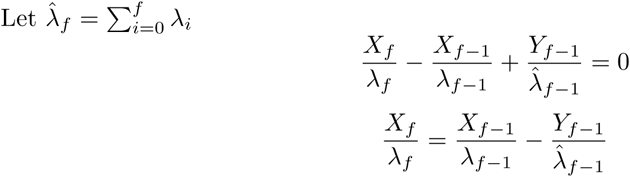

Assume that 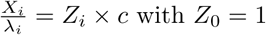 with *Z*_0_ = 1,

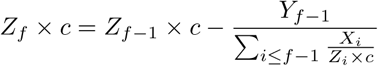

Hence,

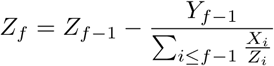

After computing *Z_f_* we can compute λ_*i*_ × *c*, and with the constraint ∑_*i*_ λ_*i*_ = 1 to get the value of

### 4.3 Read start position distribution

We have,

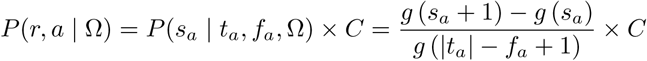

With *C* is some function that independent from *f*, hence,

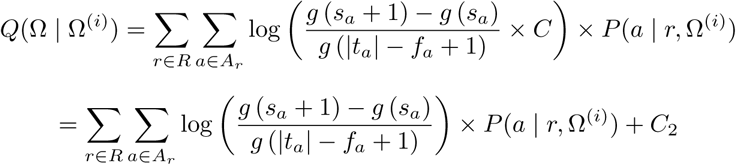

With *C*_2_ is some function independent from Ω

For simplification we assume that 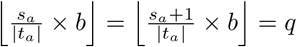 for all cases. Because transcripts are usually much longer than the number of segments, only a small gradient comes from other cases.

Therefore 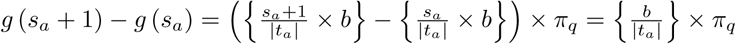

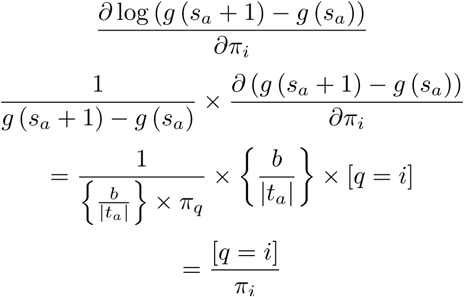

Similarly, let 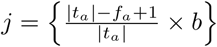, and 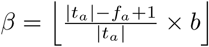

We have, 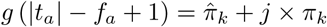

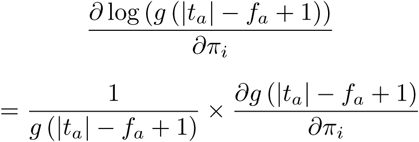

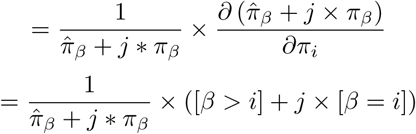

System of equation, with *α* is the Lagrange multiplier.

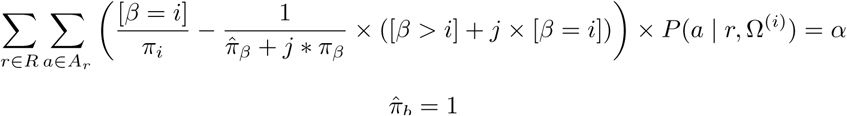

Let *X_k_* = *∑_r∈R_∑_a∈A_r,k=i__ P*(*a* | *r*, Ω^(*i*)^)

Take the difference of the equations of *π*_*i*+_1 and *π*_i_, we have,

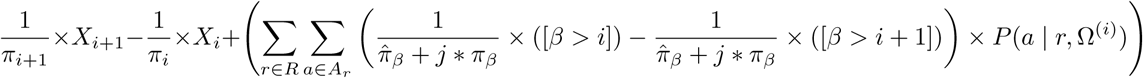

Hence,

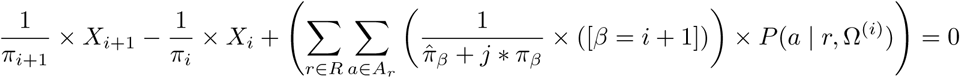

We can’t solve this equation explicitly so, we use an iterative method.

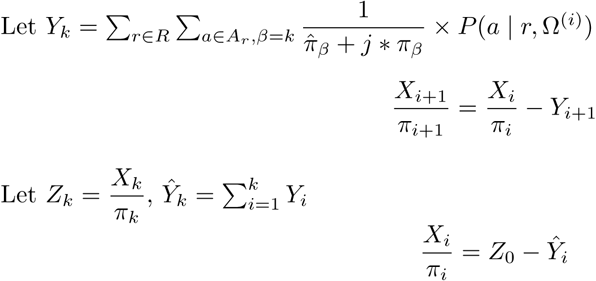

We can’t eliminate the variable *π_i_* in 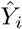 so we use an approximation from the previous value of *π_i_*. We can easily compute the estimation of *Y_i_* alongside with the EM algorithm. However there is still an unsolved variable *Z*_0_, we can use Newton method to solve the equation 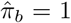 to get the value of *Z*_0_ We have,

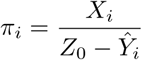

Hence,

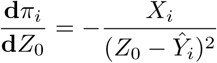

Therefore, we have the update to get *Z*_0_,

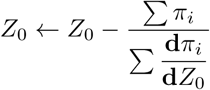

### 4.4 Orientation distribution

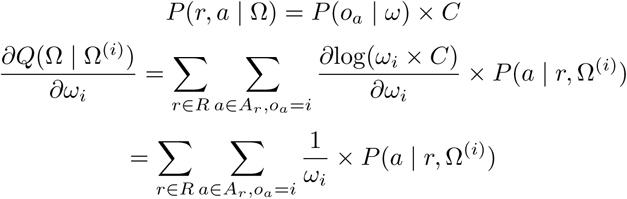

Solve the system of Lagrange equations,

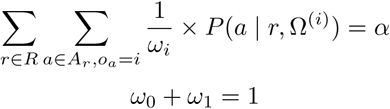

To get result,

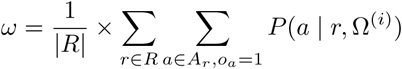

### 4.5 Alignment distribution

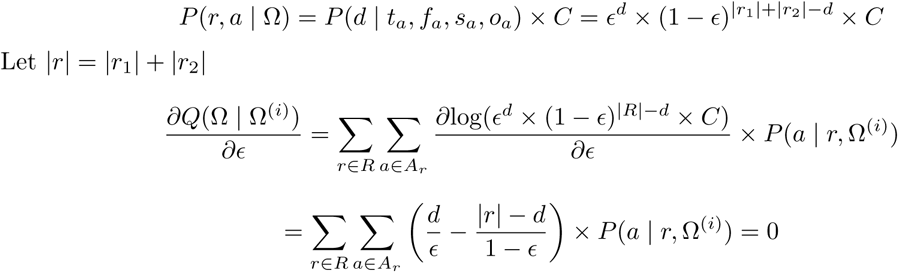

Solve the equation to get the result,

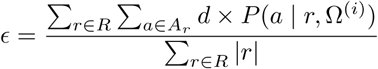

## 5 Future work

As many works have shown that variational inference is arguably a better solution for transcript abundance quantification compared to the maximum likelihood estimation. We are considering to implement a similar Hera-VB algorithm for the RSEM generative model.

## 6 Acknowledgements

This work was supported by BioTuring Inc. We would like to thank the organizer of SMC DREAM challenge for providing the simulated data sets. We also would like to thank the reviewers for reading this paper and giving us critical comments. We are in debt to Rob Patro for many insightful comments, and for very interesting conversations, so much so that Son Pham missed the flight while standing at the boarding gate.

## Footnotes

1 We also noticed that IsoEM [8] uses a different equivalent relation definition to build equivalent classes, which also requires the conditional probabilities of the alignments to be the same. Therefore, no information will be lost. This technique can be applied to our method as well. However, we think it is not a very significant optimization in our case as the number of equivalent classes would not be significantly reduced.

2 When feeding with full alignment information, Salmon [10,17] uses those data in the online phase and then the mapping conditional probabilities of each transcript are averaged for each equivalent class. While some information is still being lost due to averaging, it is less significant than other tools

3 We decided to use a “spatially-homogeneous” model for sequencing error, specifically, we only use the edit distance of the alignment and ignore the read quality and the position of those error bases, as the accuracy loss due to this simplification is minimal, while gaining a huge computational advantage.

4 Reads that map to the same set of transcripts

## References

[1] Kendall E Atkinson. An introduction to numerical analysis. John Wiley & Sons, 2008.

[2] Nicolas L Bray, Harold Pimentel, Páll Melsted, and Lior Pachter. Near-optimal probabilistic rna-seq quantification. Nature biotechnology, 34(5):525, 2016.

[3] Gregory W Corder and Dale I Foreman. Nonparametric statistics: A step-by-step approach. John Wiley & Sons, 2014.

[4] Jonathan Houseley and David Tollervey. The many pathways of rna degradation. Cell, 136(4):763–776, 2009.

[5] Ben Langmead and Steven L Salzberg. Fast gapped-read alignment with bowtie 2. Nature methods, 9(4):357, 2012.

[6] Bo Li and Colin N Dewey. Rsem: accurate transcript quantification from rna-seq data with or without a reference genome. BMC bioinformatics, 12(1):323, 2011.

[7] Naoki Nariai, Kaname Kojima, Takahiro Mimori, Yukuto Sato, Yosuke Kawai, Yumi Yamaguchi-Kabata, and Masao Nagasaki. Tigar2: sensitive and accurate estimation of transcript isoform expression with longer rna-seq reads. BMC genomics, 15(10):S5, 2014.

[8] Marius Nicolae, Serghei Mangul, Ion I Măndoiu, and Alex Zelikovsky. Estimation of alternative splicing isoform frequencies from rna-seq data. Algorithms for molecular biology, 6(1):9, 2011.

[9] Lior Pachter. Near-optimal rna-seq quantification with kallisto. https://liorpachter.wordpress.com/2015/05/10/near-optimal-rna-seq-quantification-with-kallisto/. Accessed: 2018-11-04.

[10] Rob Patro, Geet Duggal, Michael I Love, Rafael A Irizarry, and Carl Kingsford. Salmon provides fast and bias-aware quantification of transcript expression. Nature methods, 14(4):417, 2017.

[11] Rob Patro, Stephen M Mount, and Carl Kingsford. Sailfish enables alignment-free isoform quantification from rna-seq reads using lightweight algorithms. Nature biotechnology, 32(5):462, 2014.

[12] Irene Gallego Romero, Athma A Pai, Jenny Tung, and Yoav Gilad. Rna-seq: impact of rna degradation on transcript quantification. BMC biology, 12(1):42, 2014.

[13] SMC-RNA. Github - sage-bionetworks/smc-rna-challenge. https://github.com/Sage-Bionetworks/SMC-RNA-Challenge. Accessed: 2018-11-04.

[14] SMC-RNA. The icgc-tcga dream somatic mutation calling - rna challenge. https://www.synapse.org/#!Synapse:syn2813589/wiki/401442. Accessed: 2018-11-04.

[15] SMC-RNA. The icgc-tcga dream somatic mutation calling - rna challenge. https://github.com/Sage-Bionetworks/rnaseqSim. Accessed: 2018-11-04.

[16] Ravi Varadhan and Christophe Roland. Simple and globally convergent methods for accelerating the convergence of any em algorithm. Scandinavian Journal of Statistics, 35(2):335–353, 2008.

[17] Mohsen Zakeri, Avi Srivastava, Fatemeh Almodaresi, and Rob Patro. Improved data-driven likelihood factorizations for transcript abundance estimation. Bioinformatics, 33(14):i142–i151, 2017.

